# BTR: A Bioinformatics Tool Recommendation System

**DOI:** 10.1101/2023.10.13.562252

**Authors:** Ryan Green, Xufeng Qu, Jinze Liu, Tingting Yu

**Affiliations:** Department of Computer Science, University of Cincinnati, Cincinnati, 45219, USA; Department of Biostatistics, Virginia Commonwealth University, Richmond, 23284, USA; Department of Computer Science and Engineering, University of Connecticut, Storrs, 06269, USA

## Abstract

**Motivation:** The rapid expansion of Bioinformatics research has resulted in a vast array of computational tools utilized in the development of scientific analysis pipelines. However, constructing these pipelines is a laborious and intricate task, one which demands extensive domain knowledge and careful consideration at all stages. As the Bioinformatics landscape continues to evolve, researchers, both novice and expert, may find themselves overwhelmed when working in unfamiliar fields. Consequently, this may result in the selection of unsuitable or suboptimal tools during workflow development.

**Results:** In this paper, we propose the Bioinformatics Tool Recommendation system (BTR), an innovative deep learning model designed to recommend the most suitable tools for a given workflow-in-progress. BTR utilizes recent advances in graph neural network technology and introduces a novel approach, representing the entire workflow as a graph to capture essential context and structural information. Additionally, natural language processing techniques are integrated to enhance the quality of tool recommendations by analyzing associated tool descriptions. Experiments demonstrate that BTR outperforms the existing Galaxy tool recommendation system, highlighting its potential to greatly facilitate scientific workflow construction.

**Availability and implementation:** The Python source code is available at https://github.com/ryangreenj/bioinformatics_tool_recommendation

## 1 Introduction

Bioinformatics researchers utilize computational components to enable analysis and interpretation of large, complex biological data. To support this work, the practice of creating reproducible, reusable, scalable, and shareable (Wratten *et al*., 2021) analysis pipelines or computational workflows has been adopted. Several systems and standards have emerged over the years to streamline the workflow creation process, making it more accessible for individuals who may lack the technical expertise to weave workflows from scratch. Some popular workflow management systems include NextFlow (Tommaso *et al*., 2017), Common Workflow Language (Crusoe *et al*., 2022), Snakemake (Mölder *et al*., 2021), and Galaxy (Afgan *et al*., 2018). The systems are proposed to ease the creation of workflows in various ways - such as pulling from a shared *toolbox* of existing functions, automatically optimizing resource use and configurations for effective processing, handling installation and setup, and resolving any versioning or dependency issues for the user often through the use of pre-configured virtual environments like Anaconda (Anaconda, 2016) and Docker (Merkel, 2014).

Development of new workflows can be a challenging task that requires a thorough understanding of tools in the specific domain and how they interact with each other at different stages. Experienced bioinformaticians may already have the domain knowledge and coding expertise to be well-versed in their pipeline compositions. On the other hand, newer researchers, especially those with less computational backgrounds, may be more reliant on finding and using existing tools and techniques. There is a need to understand what tools are available to accomplish which tasks, how they work, and how they can be integrated with previous steps in a workflow. Finding this information can be time consuming and repetitive, requiring hours of web-surfing with abstract search concepts to find a desired function. The rapid growth of Bioinformatics has expanded the already vast catalogue of available tools, further complicating tool selection. Taking Galaxy as an example, there exists over 9,200 tools in the system with the average workflow consisting of 13 tool steps. Galaxy Toolbox saw 53% growth from 2016 to 2018 (Afgan *et al*., 2018), showing a rough 40% to 50% growth every two years since. The need to reuse functions to capitalize on the growth in Bioinformatics is complicated by this influx of new information. Discovering and integrating new tools into workflows can furthermore be hindered due to lack of training material.

It is impractical for human researchers to be knowledgeable of the complete tool catalogue. Automated solutions can assist in tool selection. Various methods have been proposed to aid in the construction of analysis pipelines. EDAM (Ison *et al*., 2013) is a Bioinformatics ontology that enables consistent annotation of entities. It is implemented in a community-sourced indexing of available tools, bio.tools (Ison *et al*., 2015), for standardized querying and has seen considerable participation. At present, bio.tools indexes 28,000+ tools, posing challenges that querying alone cannot solve, like identifying the most suitable tool that is also compatible with an existing workflow. The Automated Pipeline Explorer (APE) (Kasalica and Lamprecht, 2020) uses search constraints over bio.tools to produce abstract possibilities for workflows. While this could be particularly useful in brainstorming, the workflows must be manually validated and constructed. Workflow INstance Generation and Selection, or WINGS (Gil *et al*., 2011a,b) is a system that automatically finds implementations of abstract workflow concepts by applying specific tools. Constructing the high level workflows nevertheless requires the knowledge in choosing correct operations to perform the analysis. Kumar *et al*. (2021) propose the currently-implemented Galaxy tool recommender system, aiming to suggest downstream compatible tools following an input of preceding tool sequence. However, this approach lacks specificity in recommendation due to potentially vast compatibility sets, limiting utility. Previous automatic workflow composition methods tend to focus on identifying specific implementations for a desired concept or defining the workflow abstractly. A system that can perform both in one step would be valuable. Furthermore, works that utilize information from a workflow-in-progress commonly look at only a single step or a linear sequence of steps, which is not fully representative of the branching and winding workflow structure seen in practice.

In this paper, we attempt to address the problem of on-demand Bioinformatics tool recommendation during workflow realization and construction by proposing the **B**ioinformatics **T**ool **R**ecommendation system (BTR). We model workflow construction as a session-based recommendation (Li *et al*., 2017; Wu *et al*., 2019; Ma *et al*., 2020) problem and leverage emergent graph neural network technologies (Scarselli *et al*., 2009; Li *et al*., 2016) to enable a workflow graph representation that captures extensive structural context. This approach represents the workflow as a directed graph. A variant of the system is constrained to employ linear sequence representations for the purpose of comparison with other methods.

We conduct a comprehensive evaluation of BTR and its variants with two extensive Galaxy databases, each comprising over 1250+ unique tools, and a combined 7000+ workflows. Furthermore, we compare BTR to a baseline method created to solve a similar Bioinformatics tool recommendation problem. Lastly, we explore the viability of BTR in the age of large language models. It is found that BTR demonstrates considerable performance in the direct tool recommendation problem, firmly outperforming the baseline system. We also find that large language models do not overtake the work and demonstrate degraded performance in comparison with our specialized system given the same inputs and task.

## 2 Materials and methods

### 2.1 Preliminary definitions and representations

To accurately describe the deep learning steps of our proposed tool recommendation system - BTR, we first define how individual tools and workflows are represented. BTR requires a toolbox consisting of bioinformatics tools that are the building blocks of potential workflows. Thus, a ‘tool’ is at the lowest level of abstraction obtained from a workflow. Bioinformatics libraries often contain a collection of functions, each of which can be considered a tool in our tool box. For example, the library of BEDTools (Quinlan and Hall, 2010) consists of many important functions for sequencing read and alignment file manipulation such as *annotate* and *map*. Each function is self-contained and performs an important task. In this case, each function including *bedtools_annotate* and *bedtools_map*, along with their corresponding descriptions are added as a tool to *T* . Formally, tools are represented by a ⟨*toolboxID, description*⟩ pair.

A workflow defines the execution sequence alongside input and output connection between bioinformatics tools to perform a specific bioinformatics tasks. In this application, workflows are represented following the format of Abstract Workflow Representation (AWR) which is created following graphical displays within workflow management systems (Mölder *et al*., 2021; The Galaxy Community, 2022). The AWR *W* = (*S, C*) consists of a list of steps *S* (|*S*| ≥ 1, nodes of the graph) and the connections (edges) between those steps *C*. Each step *s* ∈ *S* is an index pointing to a tool of *T* . There may be multiple invocations of a given tool throughout a workflow; duplicates are allowed. The connections *C* represent the flow of data between tools through the workflow. We note that workflows represented by the AWR hold properties of directed acyclic graphs (DAGs).

### 2.2 Problem description

In the context of machine learning, we frame the tool recommendation task as a regression problem of predicting the most likely tool for any given workflow-in-progress. Probabilities are calculated for each candidate in the toolbox *T* . The input query to BTR is an AWR of the incomplete workflow up to the desired point of recommendation, referred to as the prefix-AWR. Two architecture variants are discussed in this paper, BTR^*g*^, which operates on full workflow graph representations, and BTR^*s*^, that consumes linear tool sequences. The output of BTR is a recommended tool that can occur in the prefix-AWR after a user-defined list of preceding steps *R*. BTR assumes that output data from steps of *R* will directly feed into the recommended tool. The system produces a set of probabilities *P* for all tools in *T* from which it can choose optimal candidates. In the example of BTR from Figure 1, *T* = [“UMI-tools extract”, “RNA STAR”, “Filter BAM”, “MultiQC”, “FeatureCounts”, …], steps *S* = [0, 1, 2], connections *C* = [(0, 1), (1, 2)]. BTR can be considered a function *P* = BTR(*w* = (*S, C*), *R* = [2], *T* = *T*). This yields a possible recommendation of *T* [argmax(*P*)] = “FeatureCounts”.

**Fig. 1:**
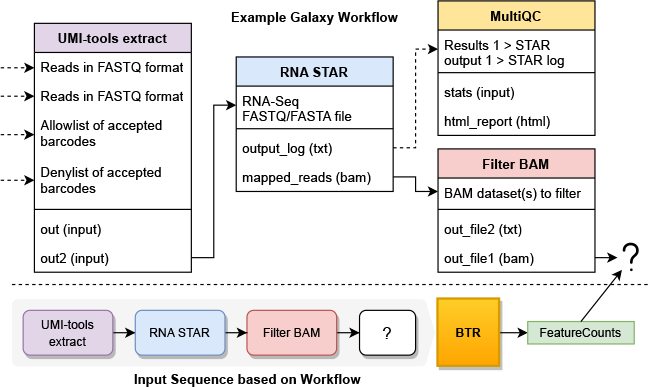
A showcase of tool recommendation by BTR. The workflow-in-progress (top) is from a tutorial^1^ about Single-Cell data pre-processing. The figure mimics how tool nodes appear within the Galaxy editor, and the connections between their inputs/outputs. In this case the user desires a tool to follow the *out_file1 (bam)* of **Filter BAM**. The next functionality from the tutorial is to create a count matrix using **UMI-tools count**, but this tool is dependent on a call to **FeatureCounts** to annotate the BAM reads with gene name. The bottom portion of the figure shows the abstraction of workflow graph/sequence to capture only tool identities and their interconnections, which serves as the input for BTR. Note that only the upstream dependencies of the tool at the desired position (denoted by ‘?’) are included. BTR correctly outputs FeatureCounts as the highest-ranked tool from 1250+ choices. ^1^https://training.galaxyproject.org/training-material/topics/single-cell/tutorials/scrna-preprocessing/tutorial.html

### 2.3 Workflow-in-progress as graph and sequence query

BTR^*g*^ is variant of the BTR architecture that employs a graph representation of a workflow-in-progress, defined as a set of upstream nodes and their connections that a recommended tool will depend on. The model internally inserts a blank query node *q*, displayed as the ‘?’ in Figure 1, and creates directed edges from user-defined preceding steps *R* to the query node. The objective is to solve the tool that replaces *q*.

Harnessing the flexibility of BTR structure, we create the second variant BTR^*s*^, where workflows are treated as linear sequences of ordered tool invocations instead of graphs. This variant is similar to the recently published approach of recommending Galaxy tools (Kumar *et al*., 2021). No query node is added to the input of BTR^*s*^ because *R*, the list of preceding steps from Section 2.2, is automatically inferred to consist solely of the most recent step in the incomplete workflow sequence.

### 2.4 Tool recommendation using graph neural networks

In this section the proposed deep learning model architecture as shown in Figure 2 is described in detail.

**Fig. 2:**
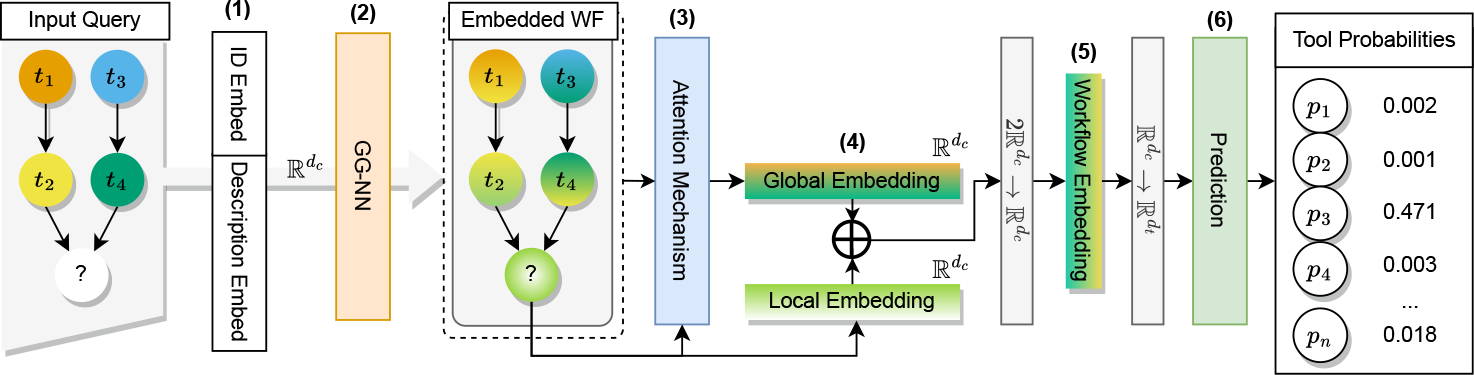
Overview of the BTR modeling architecture for Bioinformatics tool recommendation. BTR takes the input of Input Query in the format of a sequence or a graph. The prediction framework is composed of six major steps: (1) Each tool instance is encoded by an initial embedding layer (Section 2.4.1); (2) The initial embeddings continue to a Gated Graph Neural Network to learn contextural and structural features from neighboring nodes using full workflow graph; (Section 2.4.2); (3) An attention mechanism aggregates the latent graph node embeddings into a full workflow representation, which is concatenated with the representation of the last tool (4) and transformed to yield the final workflow representation vector (5) (Section 2.4.3), which is then compressed to the size of the tool embedding; (6) Tool probabilities are produced by similarity of compressed workflow representations to all tool embeddings from the toolbox (Section 2.5).

#### 2.4 Tool embedding integrating NLP description

Embeddings of toolbox ID and corresponding semantic tool description are tied together as initial node features during graph learning. Natural language processing (NLP) techniques are applied to extract latent knowledge from the semantic tool description. Including such information as a node feature allows the model to gain access to a depiction of semantic tool similarity and thus improve correlation of the usage and relationships between tools.

Tool *description* is converted to a latent vector using a sentence encoder. Sentence encoders are language models that embed sentences into 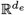 dimensional vectors that capture the semantic meaning of a sentence, useful for similarity calculation and transfer learning (Reimers and Gurevych, 2019). For this task, we use PubMedBERT (Gu *et al*., 2021), a BERT (Devlin *et al*., 2019) model that is pre-trained from scratch on a large corpora of PubMed abstracts and full-text articles. PubMedBERT embeds sentences into 768-dimensional vectors, which becomes *d*_*e*_. This encoder yields state-of-the-art results for many domain-specific NLP tasks and utilizes an in-domain vocabulary that allows tokenization of many relevant biomedical terms. We use a version of PubMedBERT^1^ that is fine-tuned for sentence embedding. This gives the representation 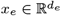 .

The index of the tool in toolbox *T* is represented by a one-hot encoded vector *t*_*ind*_ ∈ ℝ^*u*^, where *u* = |*T* | is the total number of tools in the toolbox. This vector is multiplied with a matrix of learnable weights 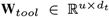 yielding a latent vector 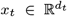, where *d*_*t*_ is a hyperparameter representing the dimensionality of the tool ID vector. **W**_*tool*_ is learned using back-propagation through time (Mozer, 1995).

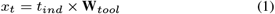

To obtain the combined encoding 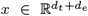 and *x*_*e*_ are concatenated. The combined hidden dimension is referred to as *d*_*c*_ = *d*_*t*_ + *d*_*e*_ throughout this section.

#### 2.4 Workflow learning using gated graph propagation

Next, workflow features and node relationships are extracted through the message passing and aggregation layer. This enables individual steps to capture both structural and feature information from their local neighbors. This capability is analogous (Hamilton, 2020) to the behavior of convolutions in convolutional neural networks (Krizhevsky *et al*., 2017). Gated graph neural networks (GG-NNs) (Li *et al*., 2016) apply the intuitions of gated recurrent units (GRU) (Cho *et al*., 2014) with the intention of better representing sequential data.

The methods of BTR are adapted for the tool recommendation problem based on the architecture of Session-based Recommendation with Graph Neural Networks (SR-GNN) by Wu *et al*. (2019). Session-based recommendation is a technique of recommender systems that only considers the recent history or “current session” of the *user* when making predictions or recommending *items* (Li *et al*., 2017; Wu *et al*., 2019; Ma *et al*., 2020). The representation of *user* is based solely on in-session data and no historical or auxiliary information is included. Our work models workflow construction as a session-recommendation task where workflows are the ‘users’ and tool steps are the ‘items’. This technique is applicable because the system should only consider the current workflow-in-progress. Wu *et al*. (2019) demonstrate how GG-NNs can be applied to the session-recommendation problem. They propose a model, Session-based Recommendation with Graph Neural Networks (SR-GNN), that uses a GG-NN layer combined with attention mechanism (Vaswani *et al*., 2017) to perform next-item prediction. The model sees improved performance over several baseline algorithms on e-commerce data, some of which utilize recurrent models themselves (Hidasi *et al*., 2015; Li *et al*., 2017).

Let matrix **A** ∈ ℝ^*n×*2*n*^ define the adjacency between nodes of the graph, the horizontal concatenation of an **A**^(*out*)^, **A**^(*in*)^ ∈ ℝ^*n×n*^, where **A**_*i*:_ ∈ ℝ^*n×*2^ are the two columns representing the out-directed and in-directed edges in **A** corresponding to a node *v*_*i*_ ∈ *V* . In this case of tool recommendation, edges are simply represented as **1** for present and **0** for nonexistent. During propagation of graph *G* = (*V, E*), the nodes *V* = [*v*_1_, …, *v*_*n*_] representations are updated by the following. Equation 2 performs message passing between the nodes of the graph using the outgoing and incoming edges defined in **A**. Here, 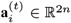 is the extracted activation of node *v*_*i*_. The remaining equations are those of GRU. Equation 3 and Equation 4 are the update and reset gates, *σ* is the sigmoid function *σ*(*x*) = 1*/*(1 + *e*^*-x*^). Equation 5 constructs a candidate state using the current state, reset gate, and previous state, where ⊙ is element-wise multiplication. Finally, Equation 6 uses the update gate to combine the previous state and candidate state to compute the final embedding. **W**_*\**_, **U**_*\**_ are learnable weight matrices and **b** is bias.

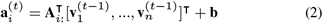

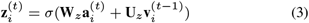

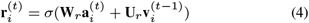

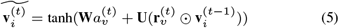

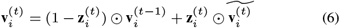

#### 2.4 Integrating local and global workflow embedding

Important contextual information is gained when looking at upstream tools in the workflow. To capture this, the intuition of Wu *et al*. (2019) is followed to model short-term and long-term preferences. A local and global workflow embedding, 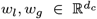, are computed and combined to aggregate the individual node embeddings into a full workflow representation. The local embedding is the representation of the most recent tool node in the workflow. We imagine there may exist rather general tools that can appear in a workflow and do not give useful contextual information; some relationships or tools in a workflow may not be as important as others. An attention mechanism is utilized to empower the model in discerning the significance of each tool in relation to others. Given the set of *n* node embeddings in a workflow, 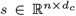, the global embedding is computed as

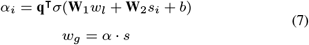

with 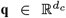, 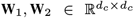and bias *b* being learnable parameters. To improve the chance of a tool being recommended that conforms with recent context, the latest tool in the sequence is emphasized by concatenating the global workflow embedding *w*_*g*_ with the local workflow embedding *w*_*l*_. The final representation is obtained by compressing this concatenation with a learned matrix 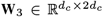and further to the space of the tool ID embedding 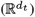 using matrix 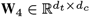

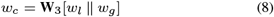

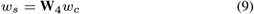

### 2.5 Tool recommendation

With the final embedded representation of a workflow-in-progress obtained, recommended tools are calculated. Each tool in the dataset is ranked based on the degree of positional similarity between its embeddings and the final workflow embedding *w*_*s*_. This is calculated simultaneously by multiplying the sequence embedding with the learned tool embedding weights matrix **W**_*tool*_ of Equation 1 as follows where *g* is a function like softmax for probabilities. *ŷ* ∈ ℝ^*u*^ is the vector of probabilities of each tool appearing next in the workflow.

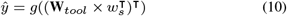

### 2.6 Model implementation

The models are implemented in Python using PyTorch (Paszke *et al*., 2019) and PyTorch Geometric (Fey and Lenssen, 2019) libraries. Each model trains to reduce the cross entropy loss between the prediction and ground truth of every workflow query, defined as

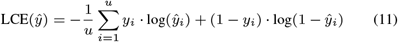

where *ŷ* are the tool probabilities from Equation 10, *y* ∈ ℝ^*u*^ is a one-hot encoded vector of the ground truth. Back-propagation through time is used to compute the gradients (Mozer, 1995). Dropout is applied to initial tool embeddings and to the aggregated node embeddings after GG-NN layer. Mini-batch Adam optimizer (Kingma and Ba, 2015) is employed to update the weights. Hyperparameters include learning rate, how much it decays by and how often, L2 penalty, batch size, number of epochs, and dropout rates. These are initialized with ranges and Bayesian optimization is used to select the values. We optimize hyperparameters over 10 training iterations and use those values for training the model.

### 2.7 Datasets and training

The proposed method is evaluated using two datasets, both which are collections of previously-created Galaxy (Afgan *et al*., 2018) workflows.

#### AllGalaxy

The first dataset, dubbed AllGalaxy, is the set of all workflows that are publicly available for viewing and download from the three UseGalaxy servers: usegalaxy.org, usegalaxy.eu, and usegalaxy.org.au (Afgan *et al*., 2018; The Galaxy Community, 2022; Afgan *et al*., 2015). Table 1 contains information about the workflow data.

**Table 1.** Information regarding the datasets collected for evaluation.

| Data Source | Workflows | Tools | Max Steps | Avg Steps | Med Steps |
| --- | --- | --- | --- | --- | --- |
| usegalaxy.org | 892 | 564 | 142 | 16.8 | 11 |
| usegalaxy.eu | 813 | 1004 | 145 | 15.1 | 10 |
| usegalaxy.org.au | 321 | 504 | 71 | 17.3 | 13 |
| Combined | 2026 | 1295 | 145 | 16.2 | 11 |
| AllGalaxy* | 1367 | 1276 | 47 | 11.8 | 9 |
| EuGalaxy* | 5782 | 1359 | 50 | 9.9 | 7 |

\* Indicates the data has been filtered to unique workflows of length 2 to 50.

#### EuGalaxy

The second dataset, EuGalaxy, is provided by Kumar *et al*. (2021) alongside their proposed Galaxy tool recommendation method. This dataset consists of public and private, potentially invalid or deleted workflows from the usegalaxy.eu server. We use a snapshot of this dataset from April 2020 provided on their GitHub repository^2^ and remove workflows labeled as erroneous.

Before training, the datasets are filtered to prevent bias and overfitting. Workflows exceeding 50 steps are discarded due to their rare occurrences. Workflows with identical tool invocations are de-duplicated to retain one instance. The filtered datasets are divided into 80% train, 10% validation, and 10% test sets, which are observed to be common split sizes in machine learning. The division occurs over the set of workflows so that individual workflows are fully contained within their respective set. Each full workflow is then iteratively split into prefix-graphs or prefix-sequences from start to finish so the model can perform tool recommendation at any stage of workflow development. These become the final queries used for training/testing. An ensemble of 20 evaluation models per variant are trained over different random train/test/val splits and the results are averaged. Optimizing hyperparameters and training all 20 evaluation models takes no longer than 8 hours for the slowest variant using an RTX 3060 laptop GPU. Models are < 30 MB on disk, scaling with toolbox size rather than number of workflows or training queries.

### 2.8 Evaluation details

The architecture is evaluated by training several variants with different experimental configurations and comparing the performance with metrics that aim to capture the system’s utility for workflow construction. The metrics measure recommendation accuracy, whereby a recommendation is considered correct if it matches the single ground truth tool for each query, as opposed to a list of potential relevant items that other recommendation systems may use. From this the following three metrics are used.

- **HR@1**: The rate at which the very first tool the model recommends matches the ground truth.
- **HR@3**: The rate at which any of the first three recommended tools match the ground truth.
- **MRR@5**: Mean Reciprocal Rank is a positional-aware measure of recommendation quality that penalizes the ground truth item appearing lower in the recommendation list. This is an appropriate metric as the model should recommend the correct item earlier to save time for a user inspecting the recommendations. It gives an idea of the model’s ability to highly-rank the correct tool.

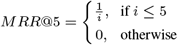

Where *i* is the 1-based index of the correct tool in the ranked tool list.

## 3 Results

### 3.1 Experiments conducted

Variants of BTR^*g*^, over graphs, and BTR^*s*^, over linear sequences, are trained and evaluated with the automated metrics from Section 2.8. We are interested in determining the impact of architecture components, so ablation studies are performed with models that build up to the full architecture. All models are prepared in the same manner as Section 2.6. The first models presented, BTR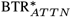, include the attention mechanism that aggregates tool embeddings into a latent workflow vector. They do not incorporate PubMedBERT description vectors and therefore only need toolbox ID as an input feature. The next models are BTR 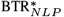, which include PubMedBERT description vectors as described in Section 2.4.1. These models do not use an attention mechanism and instead use mean-pooling to calculate the workflow node activations. The third models are BTR 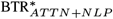 and incorporate the full BTR architecture as previously described.

BTR is evaluated against the only closely related work by Kumar *et al*. (2021), the Galaxy tool recommender (GTR) of the usegalaxy.eu server. The pre-trained model (over EuGalaxy) is obtained from the author’s GitHub repository and evaluated with the same metrics over the same test data. The pre-trained GTR model gives two sets of recommendations - one for what is defined as the high-quality, shared workflows, and another for unshared workflows. We discard the distinction between shared and unshared, the evaluation metrics are calculated for each of these sets and the higher value is taken per input to remain fair. GTR is not evaluated against AllGalaxy as we cannot obtain the unshared workflows for all usegalaxy servers that are needed to train the model.

### 3.2 Experimental findings

Results are summarized in Table 2. The key findings for the three main experiments are as follows.

**Table 2.** Performance of BTR models using evaluation metrics from Section 2.8, compared to the baseline Galaxy tool recommendation method (GTR). The values are the mean and standard deviation of the metrics from 20 evaluation models using random data splits.

| Model | AllGalaxy |  |  | EuGalaxy |  |  |
| --- | --- | --- | --- | --- | --- | --- |
|  | HR@1 | HR@3 | MRR@5 | HR@1 | HR@3 | MRR@5 |
| Baseline (Kumar et al., 2021) | - | - | - | 31.22%±2.69% | 60.46%±3.38% | 46.64%±2.76% |
| BTR<sup>g</sup><sub>ATTN</sub> | 40.70%±4.19% | 59.04%±3.86% | 50.39%±3.96% | 45.59%±1.42% | 67.66%±1.46% | 57.19%±1.32% |
| BTR<sup>g</sup><sub>NLP</sub> | 41.42%±4.44% | 59.33%±4.84% | 51.08%±4.20% | 48.22%±1.42% | 70.23%±1.40% | 59.72%±1.33% |
| BTR<sup>g</sup><sub>ATTN+NLP</sub> | **44.15%±3.78%** | **60.54%±3.89%** | **52.84%±3.68%** | **51.21%±1.33%** | **71.36%±1.32%** | **61.68%±1.21%** |
| BTR<sup>s</sup><sub>ATTN</sub> | 31.98%±3.51% | 54.28%±4.52% | 43.90%±3.79% | 38.30%±2.55% | 66.30%±3.66% | 52.93%±3.15% |
| BTR<sup>s</sup><sub>NLP</sub> | **34.43%±4.12%** | 53.55%±5.06% | 44.24%±4.28% | 44.04%±2.19% | 73.25%±3.05% | 59.23%±2.60% |
| BTR<sup>s</sup><sub>ATTN+NLP</sub> | 33.72%±4.36% | 54.27%±4.78% | 44.45%±4.41% | **47.16%±2.13%** | **75.13%±3.20%** | **61.49%±2.50%** |

#### Graph representations improve performance

BTR^*g*^ using graph representations for workflows-in-progress outperforms BTR^*s*^ using linear tool sequences. Direct, assured comparison between graph and sequence cannot be made because the data structures and training splits are different. BTR^*g*^ represents the full preceding context within a query, so there is only one query per ground-truth node in the workflows. BTR^*s*^ can have multiple queries extracted where recent tool sequences are identical, but diverge earlier upstream. Nevertheless, we perceive the performance as coverage over the datasets, for which BTR^*g*^ excels. Note the significant gap in metrics between the AllGalaxy-trained models. This suggests that the graph representation can yield strongly preferable models when less data is available. BTR^*g*^ displays improved stability across evaluation models, implying the graph representation is less sensitive to differences in data splits. The mean percentage for the metrics in the table do not show a complete picture of model performance. Figure 3 is included to visualize the evaluated metrics with different input lengths. As expected, it is observed the metrics generally increase as input length increases, with reduced effects after around length 8. The noise at larger lengths is attributed to lower numbers of available testing queries.

**Fig. 3.**
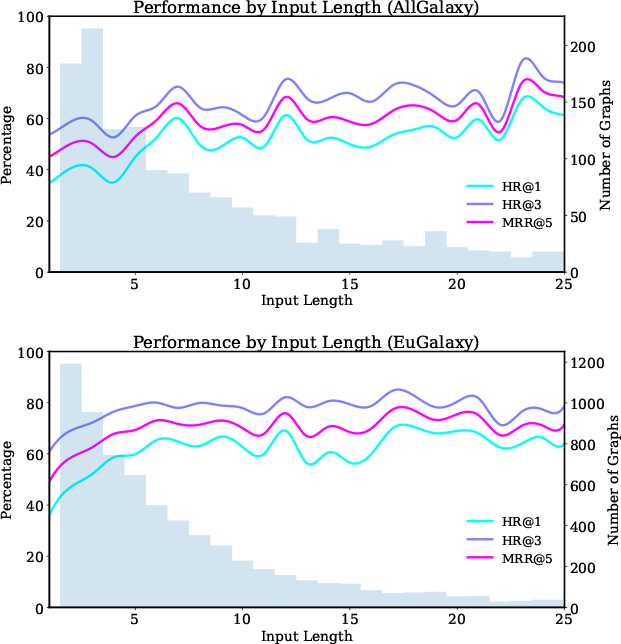
Performance of BTR^*g*^ by the input query length. The lines represent the average of the metrics from Section 2.8 at each input length and are shown as percentages matching Table 2. The bars and right-side axis show the length distribution of workflow graphs in the datasets and are included to explain noise seen in the plots.

#### Attention and NLP are impactful

The results show that both NLP and attention have notable impacts on model performance. Notice is drawn towards AllGalaxy’s BTR 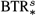 variations, which do not show as clear of a trend. We speculate that as the worst performing models, using small amounts of high-variability sequential data, it is unable to make good use of the semantic features this component provides. For the rest of the models, we observe that including short descriptions has high impact indicated by the lacking variation’s 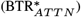 degraded performance. The attention mechanism does not have as substantial of an effect, but still a noteworthy improvement is seen from BTR 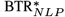 to 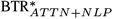. We conclude that both components are important features and in general improve the model performance.

#### BTR significantly outperforms the baseline system, GTR

BTR shows a 50% improvement over GTR, the baseline model, in recommending the correct ground-truth tool for a given query tool sequence, as measured by HR@1. The closest comparison to GTR from an architecture standpoint is with BTR^*s*^. This is because both models use linear tool sequences to represent queries and are evaluated on the exact same data and representation (comma-separated tool sequence). 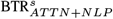 demonstrates consistent performance gain of +15% across all categories in EuGalaxy data. Note that GTR cannot run the AllGalaxy data because the unshared workflow data is not available for training. This finding underlines the potential utility of BTR during workflow construction. Furthermore, we reassert that BTR^*g*^ gives better coverage of the EuGalaxy data and an implemented system should use it to leverage the graph representation.

### 3.3 Case studies of full tool recommendation

Figure 4 shows three examples where a series of tool recommendations are conducted as a sequence, including workflows for (1) Single cell analysis, (2) COVID-19 Variation analysis, and (3) transcript assembly. The Single cell workflow demonstrates the model’s ability to chain together full workflow sequences given a starting tool. The COVID-19 workflow sees an instance where user intent could not be determined, but corrects the sequence from there after. The transcript assembly workflow shows a highly specific use case where the model cannot capture the user’s intentions without additional input. Note that the recommendations provided by our model are highly relevant nonetheless. This highlights a limitation of the model that is discussed later on.

**Fig. 4:**
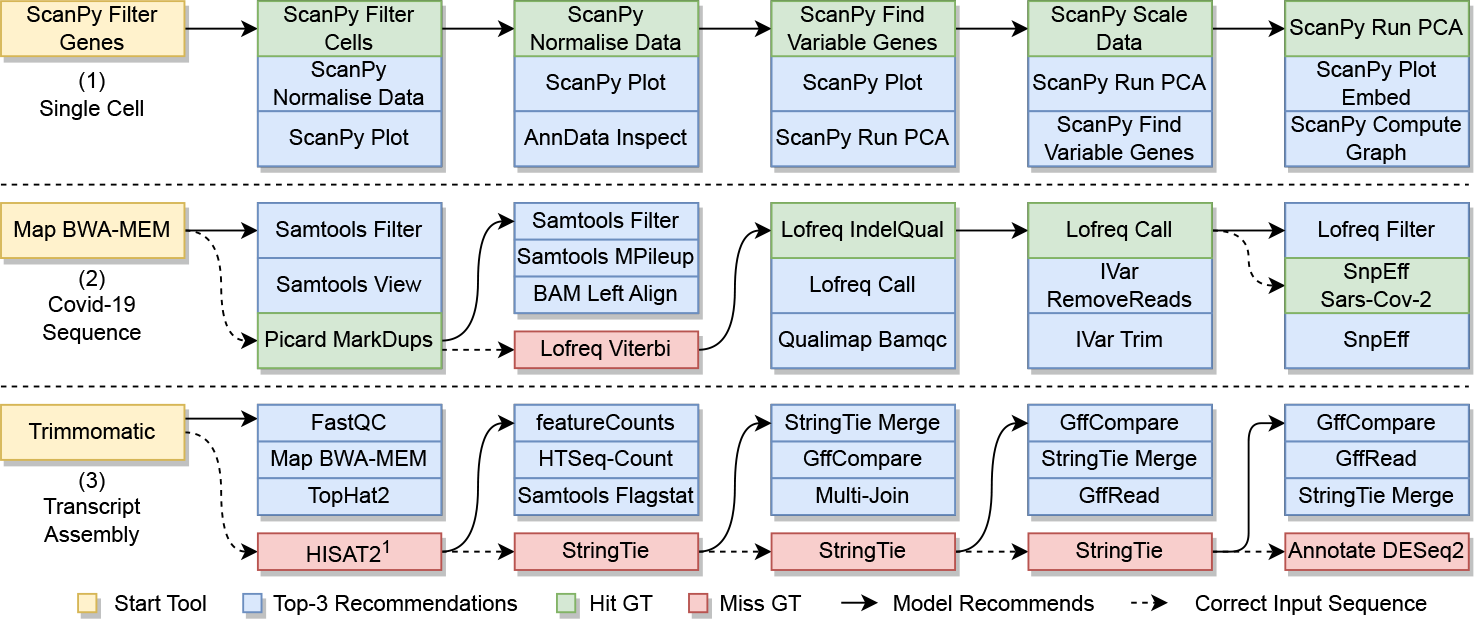
Three examples of sequential workflow tool recommendations using an AllGalaxy-trained 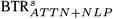 model. At each step, the correct recommendation or ground truth is included as part of the query for the next step. Examples (1) & (2) show robust utility of automated tool recommendation; Example (3) showcases highly customized workflow where recommendation accuracy can only be improved with additional user input. In cases when the recommended tools do not match ground truth, we observe that they are generally relevant and can be appropriate in different use cases. ^1^ HISAT2 is the 4^th^ ranked tool recommendation.

### 3.4 BTR comparing to large language models

AI chatbots such as ChatGPT, powered by large language models like GPT-3.5 and GPT-4 (OpenAI, 2022), are up-and-coming technologies that have the potential to be disruptive. Scientists from diverse disciplines have begun to investigate how the technology can be leveraged, including applications to Bioinformatics. Shue *et al*. (2023) provide ways ChatGPT can be used by students for solving problems and resolving errors. They find the chatbot demonstrates promising utility, however when presented with complex tasks it can start to hallucinate. In Lubiana *et al*. (2023), tips from different categories are given to describe ways this technology may enhance the routine work of Bioinformatics researchers.

We are interested in exploring how BTR stacks up with the capabilities of ChatGPT. A brief study is conducted to provide some outlook. 100 sequence queries of length 3 to 10 are randomly selected from the test set of an EuGalaxy BTR^*s*^ evaluation model. The top-3 recommendations are obtained from the model and from ChatGPT, which is constrained to the input/output format of BTR. Few-shot prompting is used to obtain the ChatGPT recommendations. Three examples of inputs and corresponding top-3 recommendations are provided, the chatbot is asked to give the top-3 tools for the new sequence in the same manner. The prompt used and 100 sample results are available as supplementary material.

The metrics for the sequences are calculated and displayed in Table 3. From the metrics alone, it appears that the general chatbot cannot perform as well as a specialized system for direct Galaxy tool recommendation. We imagine a major reason being ChatGPT is not fine-tuned for the Galaxy tool recommendation task; though it did have the potential to train on all of the tools and workflows present. In general, ChatGPT often fails to provide the desired functionality based on the workflow-in-progress. When given the correct tool for a sequence and asked why it could not provide it, the chatbot responded “… I’m not aware of the specific context or requirements of your analysis, and my response was based solely on the tool sequence you provided…” The chatbot needs more information to provide correct recommendations, which is not required by BTR.

**Table 3.** Performance of ChatGPT over 100 random samples from EuGalaxy.

| Model | HR@1 | HR@3 |
| --- | --- | --- |
| ChatGPT | 5% | 10% |
| BTR<sup>s</sup><sub>ATTN+NLP</sub> | 55% | 78% |

Table 4 includes examples chosen to highlight the behaviors of the two methods. Each row of the table contains a Sample ID corresponding to a row in the full results from supplementary material. Sample 11 demonstrates a positive result where ChatGPT ranks the correct tool in the first position. Sample 3 shows an incorrect recommendation in which BTR succeeds. Sample 31 shows ChatGPT recommending an outdated implementation of a functionality. The new tool description notes it is rewritten in modern Python, leading one to believe it is a better choice to use. The new version came out before ChatGPT’s training cut-off. In sample 56 we observe the chatbot’s tendency to hallucinate and make up tools/functionalities. The first recommended tool does not actually exist, which could mislead a user. Other cases of hallucination include ChatGPT recommending correct functionality but a partially invalid tool ID (join1 -> join2). We consider this a correct recommendation, though automated toolbox retrieval may fail in this instance. Samples 97 and 85 denote examples where ChatGPT fails to provide any recommendations. In the first case, it erroneously claims that the input tools do not exist within Galaxy. In the second, the first input tool is custom-uploaded, though the rest are available.

**Table 4.**
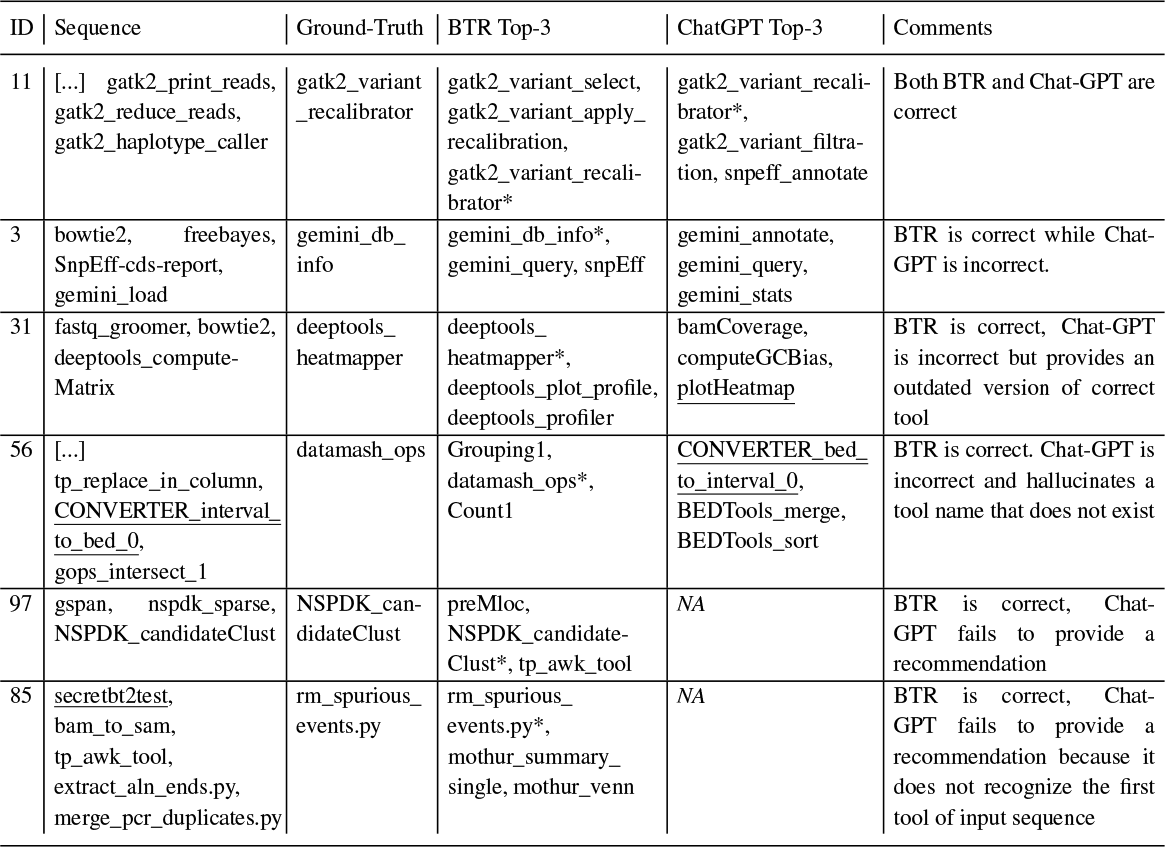
Examples of BTR recommendations alongside few-shot-prompting of ChatGPT to highlight different behaviors. The IDs match the index of the sample in the complete set. The tools shown are the tool identifiers within Galaxy, which the models take as input. An asterisk in the Top-3 highlights a correct recommendation. Underlined tools are referred to in the comments.

## 4 Discussion

Developing Bioinformatics workflows is a demanding and time-intensive endeavor, involving numerous critical factors to consider. The vast tool catalogue can add complexity to the selection process, adding further intricacies to the task. We present BTR, a novel approach to the Bioinformatics workflow tool recommendation problem. Workflow construction is framed as a session-recommendation problem and relevant techniques are applied. Recent advances in graph neural networks are utilized to capture a full workflow in a graph representation. Tool description embeddings generated by an in-domain language model are incorporated to amplify recommendation quality, along with an attention mechanism to drop weightings on low-context workflow history. Conducted experiments suggest the automatically recommended tool or tool set can provide guidance to a researcher developing a workflow, at an abstract and specific level, with only the previous steps as a query. The proposed method outperforms an existing system in Galaxy for recommending tools to construct a workflow. BTR is not overshadowed by large language models like ChatGPT. In a brief study, BTR significantly outperformed ChatGPT using few-shot prompting to recommend Galaxy tools based on workflow step sequence.

We envision the BTR architecture for workflow recommendation can be implemented as a standalone application or be incorporated into a plug-in for existing workflow management systems such as Galaxy. Its utility can be further augmented by including additional information following each recommendation, such as links to the user manual, sample codes and suggested parameterizations. We believe such system will provide instant guidance for Bioinformatics developers during the construction of workflows in an unfamiliar domain, significantly shortening development time needed. Additionally, the system has potential to enhance the quality of constructed workflows by learning from completed processes, effectively sidestepping mistakes and pitfalls.

The proposed system can be readily extended to incorporate tool parameters and configuration options. These options can be important context for tool use, as the function of some tools may change considerably with different configurations. Furthermore, configuring and optimizing the selected tools is another challenging and time-consuming task of workflow construction. Instances of a specific tool may have shared or overlapping configurations that can be matched throughout the workflows. If multiple sets of configurations with several instances each are obtained, parameter information can be incorporated into BTR. This is accomplished by extracting these configuration sets as separate tools in the toolbox. Descriptions for the different configurations could be appended with annotation data from the workflow files to further improve the recommendation quality.

Nevertheless, we foresee several actionable areas of improvements that can further increase the accuracy and utility of BTR in the next stage of its development. One limiting factor of the model performance is the amount of workflow data to train on. We speculate that the performance would continue to improve with diminishing returns as the size and generality of datasets increase. The graph representation (AWR) adopted by our model can be extended by including other workflow databases or sources including Snakemake (Mölder *et al*., 2021) and Common Workflow Language (Crusoe *et al*., 2022), both of which consist of workflow files that can be parsed into a DAG of labeled steps. The challenge with utilizing these different workflows is that, unlike Galaxy, the frameworks often do not pull from a consistent repository of tools. Consideration is needed to match tools between workflows that may have inconsistent annotations, and to further obtain corresponding tool descriptions.

Expanding user control during the recommendation process is another area for improvement. Users of the system have limited ways to control recommendations aside from the set of nodes selected to lead into the recommended tool. In the case of Figure 4 (3), additional user input at each stage such as “transcript assembly” may lead to correct recommendation. We believe the NLP component enables a modified version of the model to be prompted with natural language descriptions of the desired functionality or goal to appear next. Abstract keywords capturing intent may be used to direct the recommendations. One way to experiment with this is to shift the description embeddings to the prior step in the queries, pooling them for nodes with multiple outgoing edges. Into the last node(s), the embedded user-input text would be placed. Another option is to add the text as the description of the query node. Careful consideration needs to be taken when training and evaluating such a model, as the ground truth may be part of the input in some way.

## Supporting information

ChatGPT Behaviors Definitions

ChatGPT Few Shot Prompt

ChatGPT Sample Results

## Data availability

Instructions for obtaining and processing the datasets used to evaluate the models are available at https://github.com/ryangreenj/bioinformatics_tool_recommendation.

## Funding

The work proposed was partially funded by the University of Cincinnati Startup Grant and NSF Award CCF-2152340.

## Footnotes

1 https://huggingface.co/pritamdeka/PubMedBERT-mnli-snli-scinli-scitail-mednli-stsb

2 https://github.com/anuprulez/galaxy_tool_recommendation

## Notes

### Competing Interest Statement

The authors have declared no competing interest.

