## Supplementary material for "BTR: A Bioinformatics Tool Recommendation System": ChatGPT Few Shot Prompt

I am constructing bioinformatics analysis workflows in Galaxy. I will give a sequence of tools and want to learn tools that can come next. Please follow the format of the below examples.

Give me three Galaxy tools that can come next in this workflow tool sequence: trimmomatic -> minimap2 -> miniasm -> gfa_to_fa -> minimap2:

racon, minimap2, bandage_image

Give me three Galaxy tools that can come next in this workflow tool sequence: scanpy_filter_genes -> scanpy_filter_cells -> scanpy_filter_cells -> scanpy_filter_cells:

scanpy_normalise_data, scanpy_plot, anndata_inspect

Give me three Galaxy tools that can come next in this workflow tool sequence: fasterq_dump -> fastp -> bwa_mem -> picard_MarkDuplicates -> lofreq_viterbi:

lofreq_indelqual, lofreq_call, qualimap_bamqc

Give me three Galaxy tools that can come next in this workflow tool sequence: 
